# Fertilization reduces aphid performance but does not alter competitive exclusion between specialist and generalist species

**DOI:** 10.1101/2025.06.30.662318

**Authors:** Ayako Nakatani, Fumika Shindoh, Tatsuya Saga

## Abstract

Understanding how resource availability shapes herbivore competitive interactions is crucial for predicting pest dynamics under changing agricultural practices. Interspecific competition among herbivorous insects is mediated by host plant quality, yet mechanisms underlying competitive outcomes under varying nutrient conditions remain unclear. We investigated how fertilization affects population dynamics and competitive interactions between the legume-specialist *Megoura crassicauda* and the generalist *Aphis craccivora* on pea shoots (*Pisum sativum*) under fertilized and unfertilized conditions. Aphid population dynamics were monitored for 30 days in single-species and mixed-species treatments, and plant survival and biomass changes were assessed. Unexpectedly, fertilization reduced aphid population growth in both species, challenging conventional assumptions about nutrient effects on herbivores. Fertilization significantly affected *M. crassicauda* population growth but not that of *A. craccivora*. In mixed treatments, *A. craccivora* consistently excluded *M. crassicauda* by day 25, regardless of nutrient status. Plant mortality occurred in all aphid treatments except *M. crassicauda* under fertilized conditions. Our findings demonstrate that nutrient enrichment reduced herbivore performance but did not alleviate interspecific competition. The consistent competitive dominance of the generalist suggests that intrinsic species traits, rather than resource availability, determine competitive hierarchies. These findings have implications for pest management under changing agricultural practices.

## Introduction

In recent decades, global environmental changes such as climate change, land-use shifts, and nutrient enrichment have been recognized as major drivers of ecological processes. These changes can alter not only plant biomass and tissue structure, but also plant nutritional quality and chemical defenses. As a result, herbivorous insects that depend on these plants are affected in various ways, including distribution, feeding behavior, growth, and reproductive success (Hamann et al., 2021). Moreover, these alterations in plant condition can cascade through insect communities, influencing interactions between co-occurring herbivores.

Of particular interest is the growing recognition that plant-mediated indirect interactions among herbivores may shape competitive dynamics. Prior herbivory by one species can reduce plant quality, negatively affecting the performance of subsequently arriving species (Denno et al., 2000; Kaplan & Denno, 2007). For instance, Staley et al. (2011) demonstrated that a plant’s nutrient status could shift the outcome of interspecific competition among aphid species. However, most studies have focused on sequential rather than simultaneous interactions, and the mechanisms by which nutrient availability alters competitive outcomes remain poorly understood. In other cases, intraspecific plant interactions can alter secondary metabolite concentrations, changing herbivore preferences and distributions (Ohsaki et al., 2022). Additionally, parasitic plants can influence competitive interactions among host plants and indirectly alter insect assemblages by modifying plant defense–competition trade-offs (Shinohara et al., 2024). These findings underscore the importance of considering herbivore–plant–plant or herbivore–plant–herbivore triads, even in systems that appear to involve only two interacting species.

Understanding such interactions requires examining how resource quantity and quality shape the intensity and outcome of interspecific competition. According to classical ecological theory, competition between species may lead to either competitive exclusion or coexistence through spatial or temporal niche partitioning (Gause, 1934). However, empirical studies increasingly suggest that exclusion is not universal. Instead, coexistence is often facilitated by asymmetric resource use or environmental variability although the conditions determining competitive outcomes under varying resource availability remain difficult to predict (HilleRisLambers et al., 2012). For example, two hermit crab species in Japan have been shown to coexist by selecting different shell types (Tsuizaki et al., 2021). Similarly, Juliano (2010) demonstrated that food quality alters the relative strength of intra- and interspecific competition between two Aedes mosquito species, ultimately determining which species dominates under resource-limited conditions. Yet despite these advances, we lack a mechanistic understanding of how plant nutrient status influences the competitive balance between specialist and generalist herbivores—a knowledge gap with important implications for predicting pest dynamics under changing agricultural practices.

In this study, we focus on how changes in host plant nutrient conditions affect the strength and outcome of interspecific competition between two aphid species. Aphids (Aphididae) are phloem-feeding herbivorous insects that reproduce rapidly through parthenogenesis. At low population densities, aphids produce wingless morphs, while crowding conditions induce the development of winged individuals that disperse to new hosts. Because aphids are sensitive to both plant quality and local population density (conspecific and heterospecific), they serve as an excellent model for monitoring behavioral and population-level responses to environmental variation (Yang et al., 2021; Tena et al., 2018). Furthermore, aphids represent major agricultural pests worldwide, making an understanding of their competitive interactions directly relevant to pest management strategies.

Many studies have examined how aphid performance is influenced by plant quality. Low nutrient availability often leads to enhanced production of plant secondary metabolites and reduced nitrogen content, thereby lowering aphid fecundity and survival (Giles et al., 2002). Conversely, fertilization may promote plant growth and improve aphid performance, although in some cases competition among neighboring plants for resources may offset these benefits (Mahdavi-Arab et al., 2014). However, most research has focused on single-species responses to plant quality, with limited attention to how nutrient conditions affect interspecific interactions. Furthermore, recent evidence suggests that subordinate aphid species can detect the presence of dominant competitors and respond by reducing reproductive output or dispersing to avoid costly interactions (Li, 2019). Additional factors such as natural enemies, ambient temperature, and aphid density also modulate competitive outcomes (Petersen & Sandström, 2001; Tanaka, 1957; Tsuji & Kawada, 1985). Yet, it remains unclear how plant nutrient conditions alter the magnitude and direction of interspecific competition under shared host plant conditions.

To address this gap, we experimentally investigated how plant nutrient levels affect the competitive dynamics between *Megoura crassicauda* and *Aphis craccivora*, two aphid species that commonly co-occur on leguminous plants. Using pea shoots (*Pisum sativum*) grown under two nutrient regimes (fertilized and unfertilized), we reared the two species either separately (single-species condition) or together (mixed-species condition). This experimental design allows us to distinguish between direct effects of plant nutrition on individual species performance and indirect effects mediated through interspecific competition—a distinction critical for predicting community-level responses to environmental change.

Our primary aim was to determine whether plant nutrient enrichment promotes coexistence by alleviating competition, or conversely, intensifies competition by accelerating population growth and resource consumption. Based on resource competition theory and prior findings on plant quality effects, we developed the following testable hypotheses. (1) Under high-nutrient conditions, increased resource availability will relax interspecific competition, allowing both species to persist. This prediction follows from the stress-gradient hypothesis, which suggests that facilitation increases under benign conditions when resource limitation is reduced. (2) Under low-nutrient conditions, limited resources will intensify competition, resulting in the exclusion of one species. This expectation is based on classical competition theory, which predicts that resource scarcity strengthens competitive interactions and favor the superior competitor. We also compared mixed-and single-species treatments to assess the relative strength of intra-versus interspecific competition under varying nutrient conditions.

## Materials and Methods

### Study Organisms

This study used two aphid species from the family Aphididae: the legume specialist *Megoura crassicauda* and the generalist cowpea aphid *Aphis craccivora*. *M. crassicauda* is distinguished by its green body and dark antennae, cornicles, and cauda, and typically measures from 3.0 to 4.5 mm in length. It feeds primarily on leguminous plants, especially *Vicia faba*. *A. craccivora* has a black body and is slightly smaller (2.5 to 4.0 mm), with a wide host range within Fabaceae. In natural habitats, both species are frequently observed sharing on the same host plant. This was confirmed during field collections for this study.

### Host Plants

Commercially available pea shoots (*Pisum sativum*) were used as host plants. Approximately 60 individual shoots, each measuring 15-20 cm in height, were transplanted into plastic pots (6 cm diameter × 5.5 cm high) filled with vermiculite one week before the experiment. Plants were grown in a controlled indoor environment (20-25°C, natural light) and divided into three groups: 44 for experimental use, 10 for dry mass reference measurements, and 6 as reserve. To estimate pre-treatment biomass, 10 plants were oven-dried at 60°C for two weeks beginning three days before the experiment, and weighed using a precision balance (EK-i series, 0.01 g resolution). These values were used as baseline plant dry weights.

Aphid Collection and Inoculation

On April 29, 2024, approximately 300 wingless adults of each species were collected from a mixed-crop field in Tamatsu, Nishi-ku, Kobe, Japan (34.7°N, 135.0°E). We selected aphids based on visual assessment of vigor (active movement, intact appendages, normal coloration) and uniform body size (adult individuals of similar length within species-specific ranges). The same day, the experimental treatments were initiated by introducing aphids directly onto the host plants under laboratory conditions. Experimental Design Six treatment groups (A-F) and two control groups (G and H) were established, as summarized in Table 1. Each plant was placed in a shallow container filled with either water or a diluted fertilizer solution (see below), and positioned at least 10 cm apart to prevent cross-contamination. In mixed-species treatments, two individuals of each aphid species were introduced to each plant. In single-species treatments, four individuals of the respective species were used. Control groups received no aphids and were used to assess plant performance in the absence of herbivory.

**Table 1.** Summary of experimental conditions.

| Group | Fertilizer | Aphid species | No. of replicates | No. of aphids per plant |
| --- | --- | --- | --- | --- |
| A | No | *M. crassicauda* only | 6 | 4 (wingless adults) |
| B | No | *A. craccivora* only | 6 | 4 (wingless adults) |
| C | Yes | *M. crassicauda* only | 6 | 4 (wingless adults) |
| D | Yes | *A. craccivora* only | 6 | 4 (wingless adults) |
| E | No | Mixed (2 + 2) | 10 | 2 of each species |
| F | Yes | Mixed (2 + 2) | 10 | 2 of each species |
| G | No | None (control) | 10 | 0 |
| H | Yes | None (control) | 10 | 0 |

The fertilizer solution was prepared by diluting Hyponica liquid fertilizer (Kyowa Co., N:P:K = 5:8:12.5) to 0.2% (i.e., 500-fold dilution) in water. This solution was used in all fertilizer treatments throughout the experiment.

### Monitoring and Measurements

The experiment lasted 30 days. Aphid populations were visually counted every two days using tally counters. Water or fertilizer solution was replenished every four days to maintain consistent soil moisture levels. When winged morphs appeared, we cut their wings using fine scissors to prevent plant-to-plant migration.

After 30 days, all plants were removed from their pots, gently cleaned of soil and aphids, blotted dry, and weighed to determine fresh biomass. Each plant was then oven-dried at 60°C for two weeks and weighed again to determine final dry biomass. The duration of plant survival (in days) was recorded for each plant.

### Statistical Analysis

All statistical analyses were conducted using R version 4.5.0 (R Core Team, 2025) with the following packages: tidyverse, survival, survminer, ggplot2, dplyr, car, emmeans, and multcomp. Three datasets were analyzed: (1) aphid population trajectories, (2) plant survival time, and (3) change in dry biomass (post-vs. pre-experiment).

To analyze aphid population growth, quadratic regression models (y = ax² + bx + c) were fitted to time-series data for each treatment group. Treatment effects were evaluated using polynomial regression with interaction terms, and pairwise comparisons conducted using linear hypothesis testing from the car package.

To compare host plant survival times among treatment groups, Kaplan-Meier survival curves were constructed using the survival package, and differences were assessed using log-rank tests. Plants surviving to day 30 were treated as censored observations. The Benjamini-Hochberg correction was applied to adjust for multiple comparisons.

Changes in plant biomass were analyzed by subtracting pre-treatment dry mass from post-treatment dry mass. These differences were compared across treatments using ANOVA followed by Tukey’s HSD test for multiple comparisons. Statistical significance was set at p < 0.05.

## Results

We summarized the findings from three perspectives: (1) the effects of host plant nutrient conditions, (2) the outcomes of interspecific competition, and (3) host plant condition.

### Effects of host plant nutrient condition (fertilizer: yes/no)

#### Population dynamics differed between fertilizer treatments

Under single-species rearing conditions, maximum aphid population sizes were greater in unfertilized treatments compared to fertilized ones (*M. crassicauda*: unfertilized 77.2 ± 28.6 vs. fertilized 50.7 ± 25.2 (A vs. C: F = 17.53, p < 0.001); *A. craccivora*: unfertilized 187.1 ± 48.0 vs. fertilized 141.6 ± 43.0 (B vs. D: F = 4.00, p = 0.009)). Similarly, cumulative mean population sizes over the experimental period were higher in the unfertilized conditions (*M. crassicauda*: unfertilized 578 ± 196 vs. fertilized 292 ± 141; *A. craccivora*: unfertilized 985 ± 230 vs. fertilized 726 ± 257). The mean daily number of aphids per plant under single-species rearing conditions is shown in Figure 1 (panels I, II).

**Figure 1.**
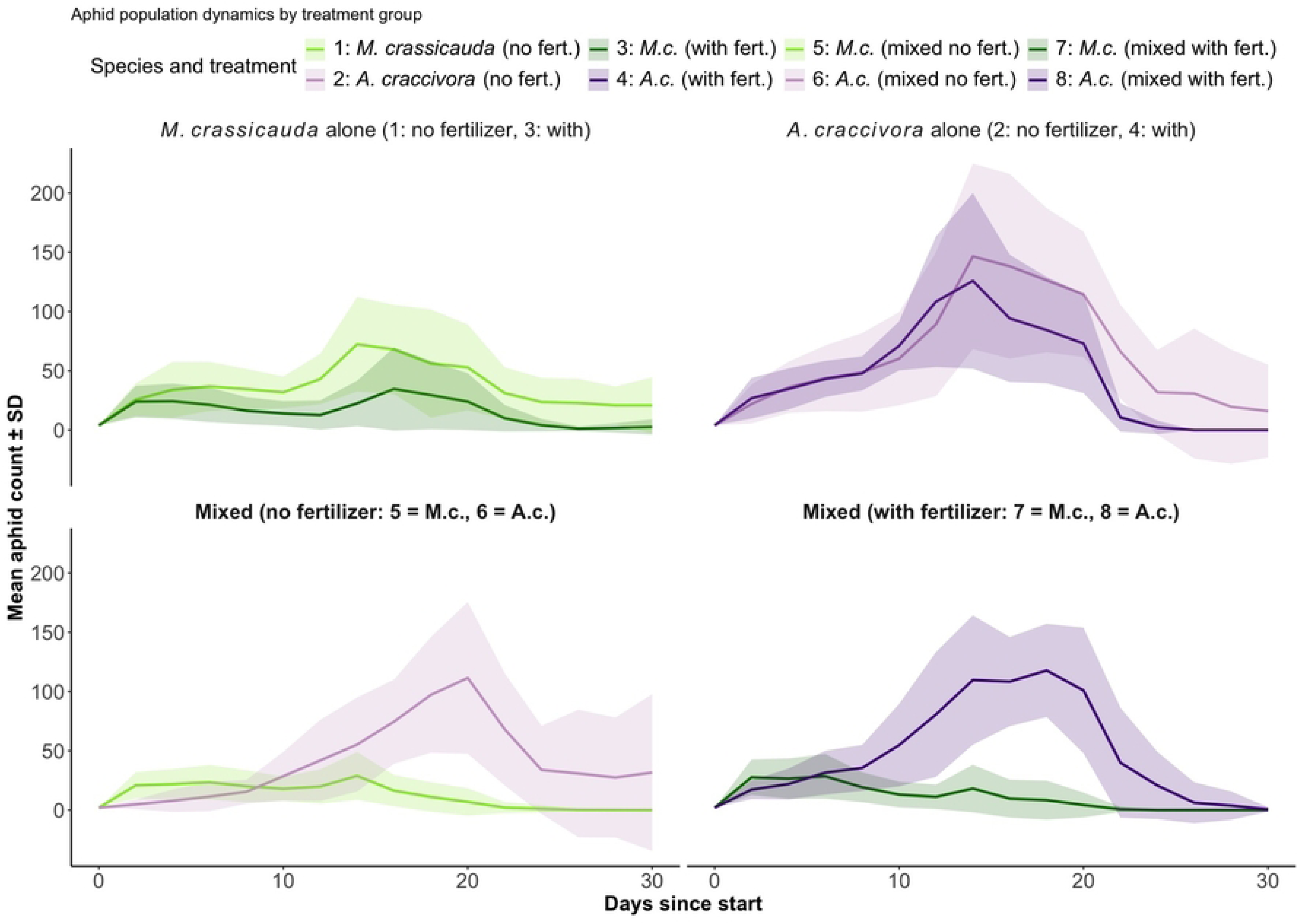
Aphid population dynamics over time under different treatments. Mean ± standard deviation (SD) of aphid counts per plant are shown over the experimental period for each treatment group. Solid lines indicate *Megoura crassicauda* and *Aphis craccivora* reared either alone (Groups 1–4) or in mixed treatments (Groups 5–8) under fertilized or unfertilized conditions. Shaded ribbons represent ±1 SD.

**Figure 2.**
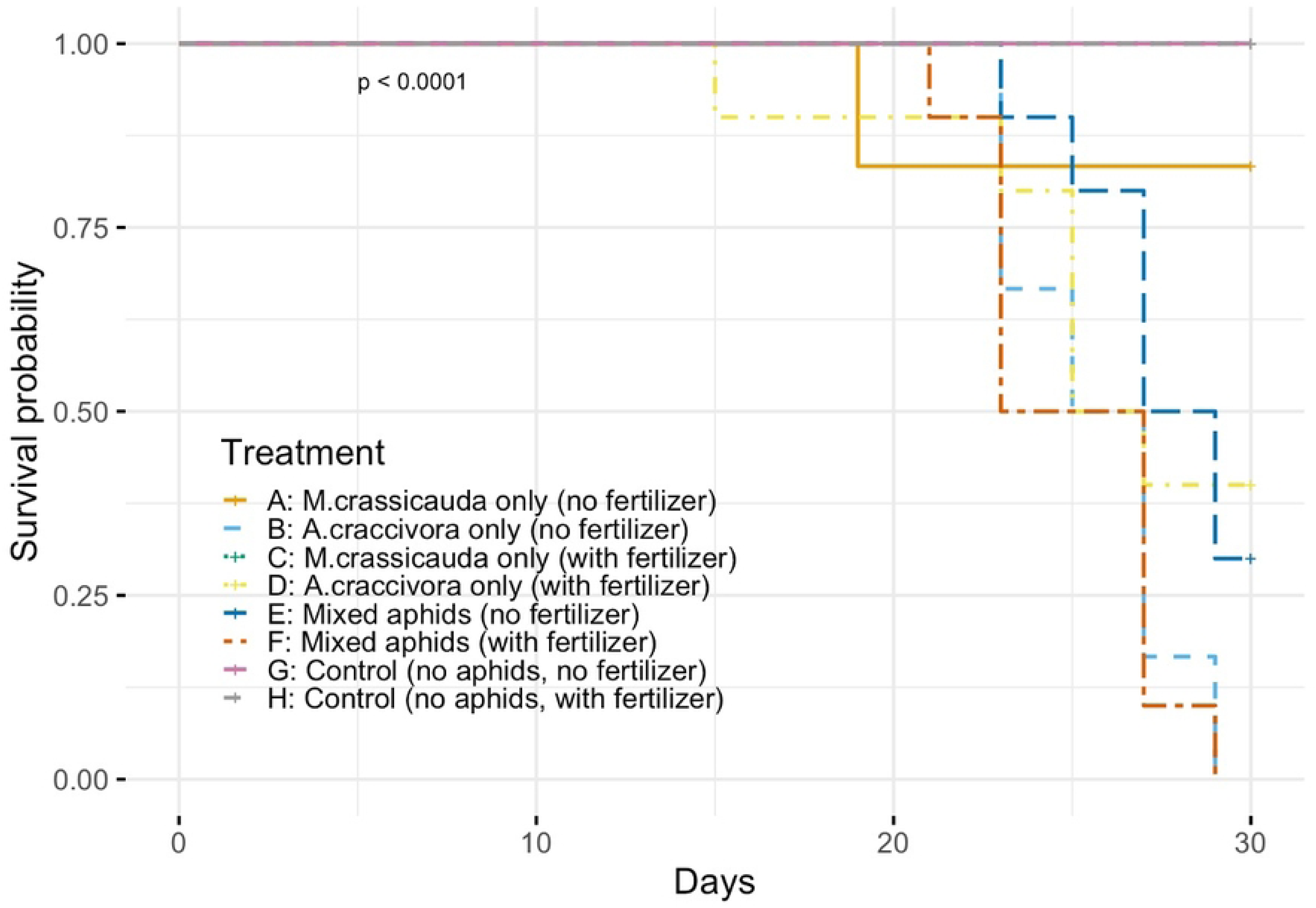
Survival analysis of pea plants infested by different aphid treatments. Kaplan–Meier survival curves show the probability of plant survival over 30 days under different aphid infestation conditions. Groups A–F represent plants infested with either *M. crassicauda*, *A. craccivora*, or both, under fertilized or unfertilized conditions. Groups G and H are uninfested control groups. Significant differences in survival time were found between treatment and control groups (log-rank test, χ²(1) = 17.2, p < 0.001). Among aphid treatments, only Group C (*M. crassicauda* with fertilizer) showed no plant mortality during the experimental period. Numbers in parentheses indicate sample sizes for each group.

Polynomial regression analysis of the relationship between the time (t) and population size (y) yielded the following model equations and associated statistics (see Figure 1-I, II, Table 2): For *M. crassicauda*, the presence of fertilizer significantly altered the population growth pattern. Both the linear (p = 0.003) and quadratic (p = 0.008) terms differed significantly between fertilized and unfertilized conditions. In contrast, no significant difference was detected for *A. craccivora* between fertilizer conditions (p = 0.229).

**Table 2.**
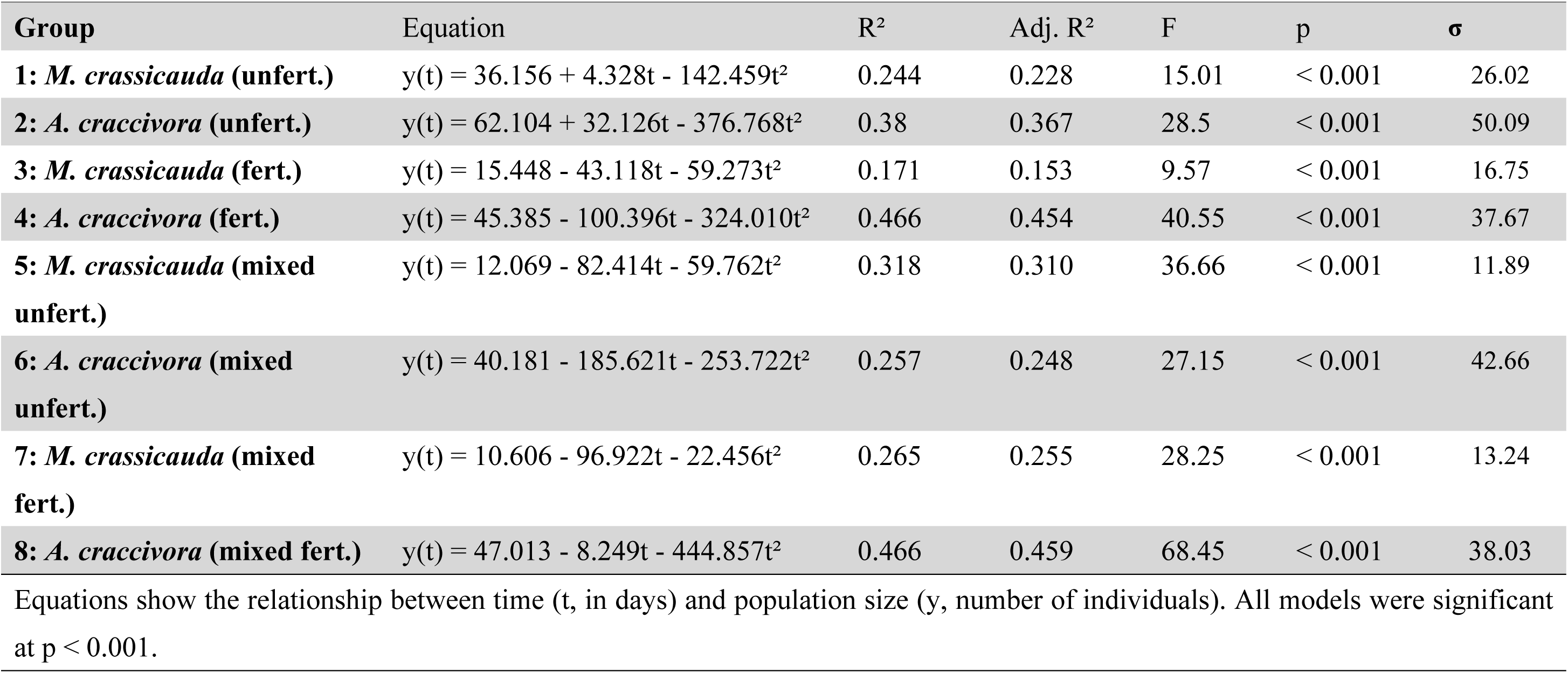
Summary of polynomial regression models for aphid population dynamics.

| Group | Equation | R<sup>2</sup> | Adj. R<sup>2</sup> | F | p | σ |
| --- | --- | --- | --- | --- | --- | --- |
| **1: *M. crassicauda* (unfert.)** | $y(t) = 36.156 + 4.328t - 142.459t^2$ | 0.244 | 0.228 | 15.01 | < 0.001 | 26.02 |
| **2: *A. craccivora* (unfert.)** | $y(t) = 62.104 + 32.126t - 376.768t^2$ | 0.38 | 0.367 | 28.5 | < 0.001 | 50.09 |
| **3: *M. crassicauda* (fert.)** | $y(t) = 15.448 - 43.118t - 59.273t^2$ | 0.171 | 0.153 | 9.57 | < 0.001 | 16.75 |
| **4: *A. craccivora* (fert.)** | $y(t) = 45.385 - 100.396t - 324.010t^2$ | 0.466 | 0.454 | 40.55 | < 0.001 | 37.67 |
| **5: *M. crassicauda* (mixed unfert.)** | $y(t) = 12.069 - 82.414t - 59.762t^2$ | 0.318 | 0.310 | 36.66 | < 0.001 | 11.89 |
| **6: *A. craccivora* (mixed unfert.)** | $y(t) = 40.181 - 185.621t - 253.722t^2$ | 0.257 | 0.248 | 27.15 | < 0.001 | 42.66 |
| **7: *M. crassicauda* (mixed fert.)** | $y(t) = 10.606 - 96.922t - 22.456t^2$ | 0.265 | 0.255 | 28.25 | < 0.001 | 13.24 |
| **8: *A. craccivora* (mixed fert.)** | $y(t) = 47.013 - 8.249t - 444.857t^2$ | 0.466 | 0.459 | 68.45 | < 0.001 | 38.03 |

Equations show the relationship between time (t, in days) and population size (y, number of individuals). All models were significant at $p < 0.001$ .

For *M. crassicauda*, population peaks were observed on days 7 (36 individuals) and 15 (68) in the unfertilized treatment, and on days 3 (22) and 17 (45) in the fertilized treatment. Both conditions showed a bimodal pattern. The second generation appeared between days 3 and 5, coinciding with the divergence in population trajectories due to nutrient treatment.

For *A. craccivora*, no substantial difference in mean population was observed between treatments up to day 10 (unfertilized: 48.6 ± 30.3; fertilized: 47.8 ± 13.1). From days 13–15, populations increased rapidly, with fertilized plants temporarily supporting more individuals (108.3 ± 50.3) on day 14. However, after day 15, the unfertilized treatment again supported higher populations, a trend that continued during the population decline phase.

#### Outcomes of interspecific competition

In mixed-species treatments, *A. craccivora* consistently dominated, with *M. crassicauda* reaching zero abundance by day 25 in all mixed treatments. Daily population monitoring revealed the following temporal pattern (Figure 1). Mean daily population sizes under mixed-species conditions are shown in Figure 1-III, IV.

In the early stage (days 1–9 without fertilizer; days 1–5 with fertilizer), *M. crassicauda* populations exceeded those of *A. craccivora* (e.g., unfertilized: 21.9 ± 12.0 vs. 7.9 ± 9.0 on day 5; fertilized: 25.3 ± 15.0 vs. 22.8 ± 11.6 on day 5). Subsequently, *A. craccivora* populations increased rapidly, surpassing *M. crassicauda* by day 11 in both fertilizer treatments (unfertilized: 28.6 ± 19.5 vs. 18.0 ± 9.7; fertilized: 56.1 ± 32.0 vs. 11.3 ± 8.9). After day 25, *M. crassicauda* was no longer observed on any plant (n = 20), whereas *A. craccivora* remained, despite a decline from its peak. Polynomial regression yielded the following equations: One-way ANOVA revealed significant differences among mixed-species treatments (Groups 5-8) (F = 63.42, p < 0.001). In both fertilizer conditions, *A. craccivora* exhibited significantly steeper increases in population than *M. crassicauda* (p < 0.001 for both linear and quadratic terms, Tukey post hoc comparisons). Single versus mixed treatment comparisons:

- A vs. B (species comparison, unfertilized): F = 12.59, p < 0.001
- C vs. D (species comparison, fertilized): F = 31.27, p < 0.001

No aggressive interactions or direct exclusion were observed. Alate morphs emerged more frequently after day 17. Colonies of *M. crassicauda* tended to form on stems, whereas *A. craccivora* colonized the undersides of leaves.

#### Host plant survival and condition

All plants in control groups (G: no aphids, no fertilizer; H: no aphids, with fertilizer) survived the full 30-day experimental period, exhibiting positive changes in biomass (G: +0.077 ± 0.083 g; H: +0.134 ± 0.089 g). In contrast, plant mortality occurred before day 30 in all aphid treatment groups except for Group C. Mean growth values were negative: −0.027 g ± 0.048 (A), −0.052 g ± 0.023 (B), −0.005 g ± 0.026 (C), −0.053 g ± 0.015 (D), −0.060 g ± 0.032 (E), and −0.105 g ± 0.032 (F).

Groups G and H are uninfested control groups. Significant differences in survival time were found between treatment and control groups (log-rank test, χ²(1) = 17.2, p < 0.001). Among aphid treatments, only Group C (*M. crassicauda* with fertilizer) showed no plant mortality during the experimental period. Numbers in parentheses indicate sample sizes for each group.

Kaplan–Meier survival analysis revealed significantly shorter plant survival time in group D compared to group C (log-rank test, p < 0.05). Furthermore, when comparing control groups (G and H) with treatment groups (A–F), plant mortality occurred significantly earlier in the treatment groups (χ² = 17.2, p < 0.001).

## Discussion

### Effects of Host Plant Nutrient Conditions on Aphid Performance

Our results demonstrate that fertilization reduced aphid population growth in both species when reared separately, contrary to the common expectation that nutrient enrichment enhances herbivore performance. This pattern was particularly pronounced for *M. crassicauda*, where fertilization significantly altered both linear and quadratic terms of population growth (p = 0.003 and p = 0.008, respectively), while *A. craccivora* showed less sensitivity to nutrient treatment.

The observed reduction in aphid performance under fertilized conditions suggests that nutrient enrichment altered plant quality in ways unfavorable to aphid development. While the specific mechanisms remain unclear, possible explanations include changes in phloem composition, leaf physical properties, or defensive compound concentrations, though plant chemical analyses would be needed to test these hypotheses. The differential response between species, with *M. crassicauda* showing greater sensitivity to fertilization, indicates that specialist and generalist herbivores may respond differently to plant quality changes. Further studies incorporating plant chemical analyses would be needed to identify the specific mechanisms involved. One possible explanation could be that fertilization altered leaf physical properties, though direct measurement would be needed to confirm this. Previous studies show this can serve as a physical defense mechanism (Hanley et al., 2007), though further investigation would be needed to confirm this mechanism in our system. However, the effectiveness of mechanical defenses varies among feeding guilds, with sap-sucking insects like aphids showing less consistent responses to leaf mechanical properties compared to chewing insects (Caldwell et al., 2016).

The differential responses between species indicates that generalist and specialist herbivores may respond differently to plant quality changes induced by fertilization. *A. craccivora*, as a generalist, may possess broader physiological tolerance to varying plant chemical compositions, while *M. crassicauda*, as a legume specialist, may be more sensitive to specific changes in its preferred host plant chemistry.

### Outcomes of Interspecific Competition

The consistent dominance of *A. craccivora* over *M. crassicauda* in mixed treatments, regardless of fertilization status, demonstrates asymmetric competition leading to competitive exclusion. This outcome occurred despite *M. crassicauda* initially establishing higher population densities in the early phase of the experiment (days 1-9), suggesting that early colonization advantage does not guarantee competitive success.

Similar patterns have been documented in ectomycorrhizal fungi, where species that gain initial competitive advantage through spore-based colonization can lose their dominance when competition shifts to mycelial-based interactions (Kennedy et al., 2011). In some host-parasitoid systems, specialists with narrow host ranges are known to excel at host detection, although generalists may gain advantage under certain conditions.

The temporal dynamics revealed distinct phases of competition: initial establishment (days 1-9), competitive reversal (days 10-25), and exclusion (day 25 onwards). Such three-phase dynamics are consistent with theoretical models of competition–colonization tradeoffs, where early colonizers may be displaced by superior competitors depending on spatial resource structure and tradeoffs between dispersal and competitive ability (Bogar & Kennedy, 2017; Smith et al., 2018). The mechanisms underlying the competitive exclusion of *M. crassicauda* by *A. craccivora* remain to be determined. Although no direct aggressive interactions were observed, several non-exclusive hypotheses could explain this pattern: (1) differential resource utilization efficiency, (2) chemical interference through honeydew or other secretions, or (3) differential responses to induced plant defenses. The spatial segregation observed (*M. crassicauda* on stems, *A. craccivora* on leaf undersides) suggests possible niche differentiation, yet this was insufficient to prevent competitive exclusion. Further research incorporating detailed behavioral observations and resource utilization measurements would help elucidate these mechanisms.

The differential impacts of the two aphid species on plant survival and growth may reflect their contrasting ecological strategies as specialist versus generalist herbivores. Generalist herbivorous insects often need to consume significantly more plant biomass when feeding on less suitable hosts in order to meet their nutritional requirements (Münzbergová & Skuhrovec, 2020). In contrast, specialist herbivores, having evolved tolerance to their host plant’s defensive compounds, may achieve sufficient nutrition with less feeding and thus cause relatively lower plant damage (Ali & Agrawal, 2012). Our findings align with recent meta-analyses showing that specialist herbivores generally cause less damage to their host plants compared to generalists (Ali & Agrawal, 2012), though the underlying mechanisms remain to be fully elucidated.

Consistent with this, in the present study, *M. crassicauda*, a specialist on fabaceous plants, caused relatively less plant mortality under fertilized conditions. In contrast, *A. craccivora*, a generalist, exerted consistently strong negative impacts on plant health irrespective of fertilization status. Differences in plant nutritional quality and defensive traits are known to significantly influence herbivore feeding strategies and performance, particularly distinguishing specialists from generalists (Kariñho-Betancourt et al., 2023).

Furthermore, the observed fertilization effects may be linked to how plant nutritional status influences aphid feeding behaviors. Enhanced plant nutrition under fertilized conditions may have allowed *M. crassicauda* to meet its nutritional requirements more efficiently, though the specific mechanisms require further investigation. Conversely, *A. craccivora*, with broader dietary tolerance, likely maintained aggressive feeding regardless of plant nutritional status. This highlights how resource availability, plant tolerance, and plant resistance interactively influence herbivore-induced plant damage (Wise & Abrahamson, 2005; Lin et al., 2021).

### Host Plant Responses and Implications

Plant survival analysis revealed contrasting impacts of the two aphid species on host plant health. The complete survival of plants infested with *M. crassicauda* under fertilized conditions (Group C) contrasts sharply with the mortality observed in other aphid treatments (e.g., Group D: *A. craccivora* with fertilizer, mean survival = 23.3 ± 3.9 days), indicating species-specific differences in plant impact intensity. Specialist herbivores have typically evolved adaptations that enable them to effectively cope with specific host plant defenses (Kariñho-Betancourt et al., 2023), allowing them to obtain necessary nutrition with relatively less plant damage. In contrast, generalist herbivores, which feed on a wider range of plant families, often require increased plant biomass consumption when confronted with highly defended plant species, as their detoxification mechanisms are less adapted to specific defenses. Indeed, previous studies comparing specialist and generalist herbivore performances on wild versus cultivated Brassica populations demonstrated that generalists caused greater plant damage than specialists due to their less specialized detoxification abilities (Gols et al., 2008).

The current study confirmed that *A. craccivora* consistently imposed greater physiological stress on host plants, evidenced by earlier plant mortality and more substantial biomass reduction. This pattern persisted across both single-species and mixed-species treatments, suggesting that *A. craccivora* depletes plant resources more rapidly and/or inflicts greater feeding damage compared to *M. crassicauda*.

The interaction between fertilization and aphid species effects on plant survival suggests complex tri-trophic relationships involving nutrients, herbivores, and plant fitness. Under fertilized conditions, plants may better tolerate the feeding pressure from specialist herbivores (*M. crassicauda*), but remain vulnerable to generalist species (*A. craccivora*).

Tolerance refers specifically to a plant’s ability to withstand or recover from herbivore injury through growth and compensatory physiological processes (Koch et al., 2016). Studies indicate that plants growing under nutrient-poor conditions generally exhibit low overall fitness but can become more tolerant to leaf herbivory due to adaptive allocation of limited resources (Wise & Abrahamson, 2005). Conversely, under nutrient-rich conditions provided by fertilization, plants may have enhanced capacity for recovery after herbivory, thus improving tolerance mechanisms. Indeed, resource limitation such as reduced water availability has been shown to lower the tolerance capacity of plants, measured by their regrowth and reproductive recovery following herbivory (Lin et al., 2021). These findings underscore how resource availability fundamentally shifts the balance between plant tolerance and resistance strategies in response to herbivory.

### Ecological and Applied Implications

These findings have important implications for understanding aphid community dynamics under changing agricultural management practices. The fertilization-mediated reduction in performance of certain aphid species (specialists) suggests that particular nutrient management strategies might unintentionally offer pest control benefits. However, this effect is species-specific, and competitive displacement toward more damaging generalist species could lead to increased crop damage.

The consistent competitive dominance of *A. craccivora*, regardless of fertilization status, indicates that environmental modifications affecting resource availability may not alter fundamental competitive hierarchies between herbivore species. Consequently, predicting pest community composition under different management scenarios remains feasible within a certain range of environmental variation.

From an agricultural perspective, the observed differential impact of the two aphid species on plant health implies that competitive displacement favoring *A. craccivora* may result in increased crop damage even if overall aphid abundance remains constant. Previous studies support the hypothesis that generalist herbivores typically cause greater feeding damage to plants than specialist herbivores, reinforcing the importance of considering herbivore identity rather than simply abundance when making pest management decisions (Ali & Agrawal, 2012; Kariñho-Betancourt et al., 2023).

Moreover, plant tolerance (capacity to recover from herbivory) and resistance (capacity to prevent herbivory) can vary depending on plant nutritional and moisture conditions (Wise & Abrahamson, 2005; Lin et al., 2021). Enhanced plant nutritional status through fertilization potentially improves plant tolerance, particularly against specialist herbivores. Thus, fertilization management may play a crucial role in enhancing plant defensive capabilities and in achieving effective pest control.

## Study Limitations and Future Directions

Several limitations should be acknowledged. First, the experiment was conducted under controlled laboratory conditions using a single host plant species and a specific fertilizer composition. Field conditions involving multiple plant species, varying environmental stresses, and complex arthropod communities may yield different competitive outcomes. Second, the absence of natural enemies in our experimental system differs from field conditions where predation and parasitism can alter competitive outcomes. Third, the mechanisms underlying the observed competitive asymmetry remain unclear from our behavioral observations alone.

While this study provides valuable insights into aphid competitive dynamics, the specific biochemical mechanisms underlying the unexpected negative effects of fertilization on aphid performance remain to be elucidated. Future investigations should incorporate: (1) comprehensive plant metabolomic analyses to understand how nutrient enrichment alters plant chemistry, (2) detailed behavioral analyses and resource utilization measurements, and (3) field studies that include natural enemy complexes to validate laboratory findings under realistic conditions.

These findings contribute to our understanding of how environmental changes may restructure herbivore communities and highlight the importance of considering competitive interactions in predicting ecological responses to anthropogenic change.

The predictable nature of competitive outcomes demonstrated here provides a foundation for developing more effective pest management strategies under changing agricultural conditions.

## Acknowledgments

The authors thank the JST Global Science Campus ROOT Program for supporting this study. The authors are also grateful to Akashi Kita High School for kindly providing laboratory equipment essential for the experiments.

